# ChemPLAN-Net: A deep learning framework to find novel inhibitor fragments for proteins

**DOI:** 10.1101/2021.08.08.455375

**Authors:** Michael A. Suarez Vasquez, Mingyi Xue, Jordy H. Lam, Eshani C. Goonetilleke, Xin Gao, Xuhui Huang

## Abstract

Fragment-based drug design plays an important role in the drug discovery process by reducing the complex small-molecule space into a more manageable fragment space. We leverage the power of deep learning to design ChemPLAN-Net; a model that incorporates the pairwise association of physicochemical features of both the protein drug targets and the inhibitor and learns from thousands of protein co-crystal structures in the PDB database to predict previously unseen inhibitor fragments. Our novel protocol handles the computationally challenging multi-label, multi-class problem, by defining a fragment database and using an iterative feature-pair binary classification approach. By training ChemPLAN-Net on available co-crystal structures of the protease protein family, excluding HIV-1 protease as a target, we are able to outperform fragment docking and recover the target’s inhibitor fragments found in co-crystal structures or identified by in-vitro cell assays.

---

Drug discovery is a long and expensive process, with the pharmaceutical industry estimated to have 2.8 billion USD in development costs and a drug target to approval timeline of around 12 years.^1^ The main challenge, given a drug target, is finding an adequate drug molecule that demonstrates selective inhibition in small concentrations. From a chemical perspective, this requires the drug molecule to interact with the binding site of the drug target, both on an atomic scale (favourable molecular force interactions) and a geometric scale (minimal steric hindrances, large topological interaction interface). The potential drug molecule space with a molecular weight less than 500 Da is estimated to be on the order of 10^60^ molecules,^2^ making a brute force search to find the ideal drug molecule for a drug target, even with modern technology, costly in terms of time and physical resources.

Computational prediction of small-molecule inhibitors based on the structure of the protein target is an important first step in the drug discovery process. However, screening heaps of small-molecules while considering the structural physicochemical properties of the target proteins is computationally challenging. Accordingly, Fragment-Based Drug Design (FBDD) has emerged as an effective strategy to overcome these challenges.^3^ Chemical fragments are light-weight organic moieties (usually ≤250 Da) that may be linked up or refined to improvise drug-sized compounds. The combinatorial combination of three or more different fragments of a 10^5^ sized fragment database can lead to 10^15^ different compounds, which is orders of magnitude larger than the upper limits of most high-throughput screening (HTS) methods at 10^6^ compounds.^4^ Predicted or observed fragment binding modes can also be used to guide compound optimization under structure-based settings using regions in the binding site inaccessible to larger molecules.^5^ Furthermore, fragments allow the exploration of chemical features at a more detailed resolution with higher specificity on local binding sites compared to larger ligands. Existing approved drugs can be easily screened against the availability of predicted fragments within their chemical structures for their repurposing on new drug targets.

A few computational tools have been prepared for the study of fragment-protein interactions.^6–8^ Most of them are inexpensive mapping algorithms coupled to a conventional docking procedure that maps fragments to their respective binding sites on proteins. For instance, the FTMap algorithm uses a sufficiently simple interaction-energy function (where fast Fourier transform can be applied) to allow rapid initial sampling of fragment-binding sites.^9^ In addition, PLIff, an inexpensive force-field derived entirely from ligand-protein contacts on the Protein Data Bank (PDB), can perform fragment-mapping as well as docking.^10^ However, despite their usefulness in detecting fragment-binding sites, these methods carry significant limitations. Since they require interaction-energy/distance calculations, the fragment mapping has to be considered one fragment-protein pair at a time, which reduces their efficiency when applied to an extensive fragment library that has to be carefully curated. Furthermore, as small polar fragments are able to bind to many polar residues on the protein that is in close proximity with marginal difference in binding energy, it is challenging to effectively isolate the contributions of individual binding sites. Workarounds include user-imposed restrictions for docking, such as limiting the search grid box for the fragments to bind on the protein. However, treating the fragment as an independent specie neglects the hierarchical connection to the original binding ligand and leads to a loss of information.

The method used in our study is based on a similar underlying assumption to the computational tools mentioned above. Namely, that similar drugs may share similar drug targets due to similar local binding environment interactions and topological fit. However, despite their proven usefulness, accessibility and transferability to other proteins is challenging as many different data streams are required. Furthermore, newly discovered drug inhibitors are often restricted to a pre-selected set of inhibitors as part of the network and do not provide the structural details of the interaction.

To this end, we built the deep learning architecture reported in this work based on data from the FragFEATURE framework.^11,12^ FragFEATURE is an instance-based algorithm that infers fragment-binding sites based on the assumption that proteins differing significantly in their active pockets could still share highly similar binding sites (local structural environments or micro-environments), and that highly similar local structural environments bind to more or less the same set of fragments. FragFEATURE is able to vectorize multiple binding environments centred around a protein residue non-carbon hetero-atom to characterize local physicochemical properties in terms of their discrete radial distribution and assign a corresponding ligand fragment label. FragFEATURE was shown to be robust in predicting fragment-binding sites shared among proteins with little structural or sequence homology. However, the original protocol is based on a nearest neighbour search, ^11^ and therefore, has limited utility as a predictive program when querying for novel fragments, as all predictions originate from existing data. We note that this does not contain any ‘learning’ process, and thus no intrinsic rules can be established that would be able to reveal higher-order relationships between different features over a wider range of protein environments. From a computer science perspective, finding the correct inhibitor fragment for a given protein structure environment using a deep learning framework is significantly more challenging. The fragment prediction classes are in the order of roughly sixty thousand fragments (multi-class), and each environment has the capability to bind to more than one fragment (multi-label) due to the chemical similarity of fragments (e.g., benzene, toluene, and xylene). Furthermore, the available ground truth data is derived from, and limited to, published experimental co-crystal structures, thus negative (non-binding) examples are not available, making verification of false positives very difficult.

Proteases are an important class of enzymes that catalyze proteolysis, the breakdown of proteins into smaller polypeptides, which is an essential step in protein catabolism, cell signalling, and viral infectious cycles.^13^ Viral proteases are enzymes encoded by the genetic material of viral pathogens to catalyse proteolysis in cellular proteins or viral polyprotein precursors. Inhibition of viral proteases is thus a pivotal step to combat new viruses as it will halt their replication process inside the infected host cell, which is particularly useful when pre-exposure vaccinations have not been developed. For example, the viral HIV-1 protease is an interesting drug target due to it being the causative agent of AIDS, a disease that infects over 40 million people worldwide.^14,15^ The development of traditional vaccines for HIV is challenging and has failed multiple times as a result of the virus’ mutative nature.^16^ The key function of HIV-protease in the virus replication cycle is the break down of precursor polyproteins into structural proteins and other essential viral enzymes such as reverse transcriptase and integrase.^17^ This critical function has made it a popular drug target in the past, with multiple known inhibitors found to date. HIV-protease is very well studied with over 100 PDB co-crystal structures publicly available and thus an ideal candidate for the real-life validation of our proposed model. There are also numerous in-vitro studies available that allow us to validate our proposed model from the experimental side.

Here we present a novel method called “Convolutional Neural Network based on Chemical features of Protein-Ligand Association” (ChemPLAN-Net) that combines the underlying data of the FragFEATURE protocol with a state-of-the-art convolutional neural network (CNN) architecture^18^ to approach multi-fragment mapping for the HIV-1 protease. The nearest-neighbour search in the FragFEATURE protocol is re-designed by a deep learning framework that learns from the FEATURE vectors and is expanded through the additional use of molecular Morgan Fingerprints. Computationally, this is a multi-label and multi-target problem for which currently there is no out-of-the-box solution. Using our biochemical insight, we develop a CNN-based classification architecture as the framework incorporating both the protein and ligand descriptors. We benchmark and validate ChemPLAN-Net using HIV-1 protease and then make a blind prediction using experimental literature as support. Firstly, we demonstrate the conceptual advancement of ChemPLAN-Net over the traditional FragFEATURE protocol by highlighting its ability to discover novel fragments not present in the binding training data. Secondly, we train our deep learning model and a machine learning implementation on all protease co-crystal structure entries except for HIV-1 protease and homologs that share ≥40% sequence similarity. Thirdly, we compare their performances against a benchmarked docking method by recovering native ligands from our HIV-1 protease validation set. We investigate the limitations and potential shortcomings regarding atom coverage when mapping the predicted fragments back onto native inhibitors, and test the assumptions of our prediction protocol. We compute the precision rate and highlight the necessity of clustering the fragment results. Finally, we compare the predicted fragment results of our framework to fragmentised proposed inhibitors in a blind prediction from the literature to evaluate their potential effectiveness.

## RESULTS AND DISCUSSION

### Overview of ChemPLAN-Net framework

ChemPLAN-Net is an end-to-end pipeline developed to train only on the three-dimensional PDB protein co-crystal structures (Fig. 1a) without requiring additional input data, such as experimental assays. The goal of ChemPLAN-Net is to reveal higher-order relationships between different physicochemical features over a wider range of protein environments and ligands, to which end, a deep learning convolutional neural network is used. The protein surface characteristics extracted with the FEATURE framework according to the training protocol (See Methods, Fig. 1b) are count-based physicochemical metrics that are rotationally invariant, and thus independent of the absolute 3D coordinates of the protein structure. This allows us to use proteins of varying sizes as training data since local environments can always be captured using the radial 6 × 80 FEATURE vector centred on the residue hetero-atoms, irrespective of protein size. The native ligands are decomposed into fragments using our curated 59,732 sized fragment database. The FEATURE environments are paired with the Morgan Fingerprints of their respective native ligand fragments and combined with an equal amount of artificially created nonbinding fragment data based on their fingerprint dissimilarity (see Methods). This joint dataset is used to train the deep learning framework (Fig. 1c). The parameters of the network are optimized through standard backpropagation using the categorical cross-entropy loss given two classes (binding and nonbinding). After the training is completed for ChemPLAN-Net, FEATURE vectors can be extracted on the surface of a query protein according to the prediction protocol (Fig. 1d). The environments of the binding pocket can be tested against all fragments in the fragment database, resulting in a sorted list of probabilities with top predictions ready to be mapped onto query inhibitors (Fig. 1e).

**FIG. 1.**
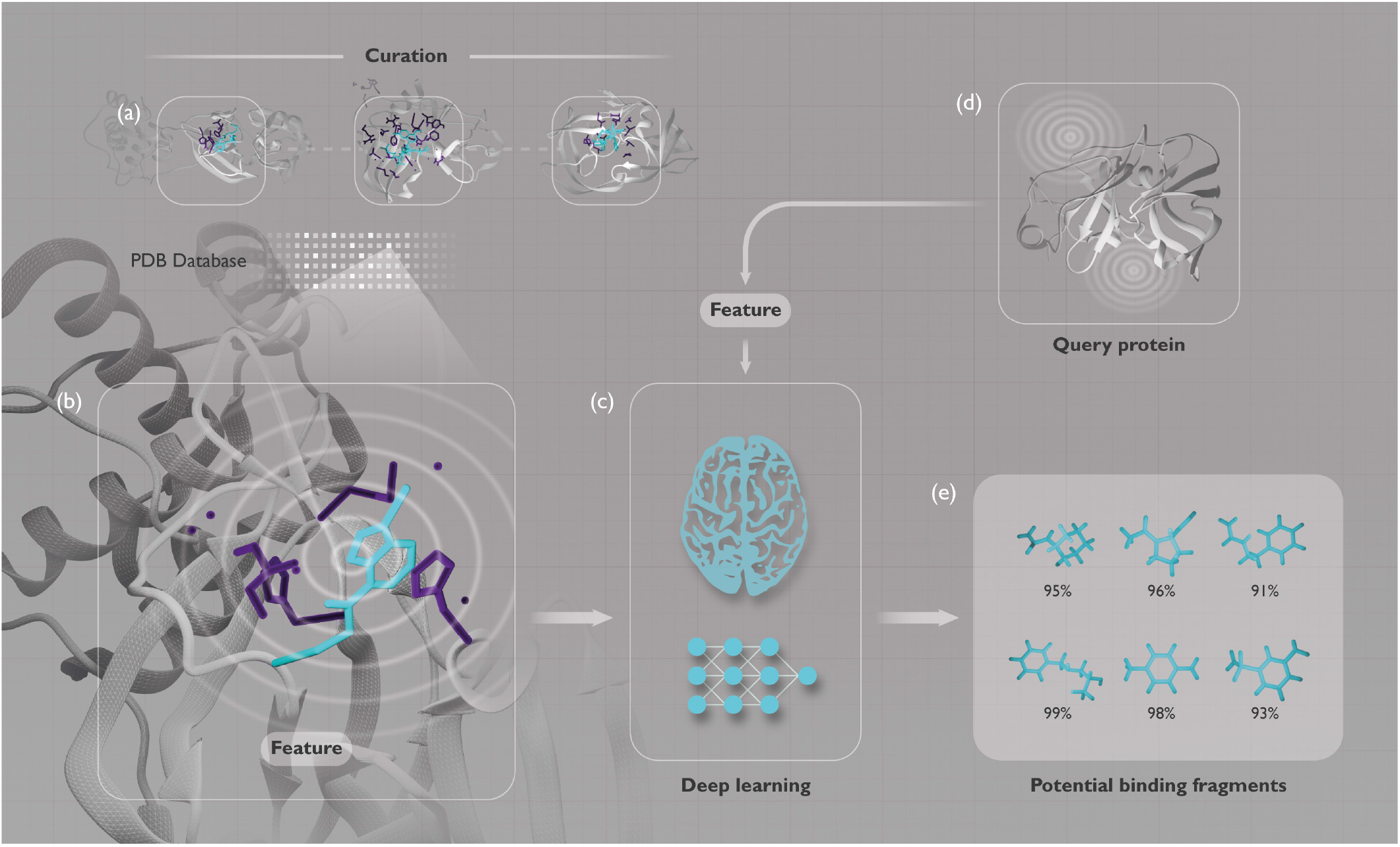
Overview of the ChemPLAN-Net pipeline. (a) Curating the 74,000 Protease co-crystal environments from the PDB database. (b) Extraction of the 1.7 M FEATURE vectors from the structure environments and the corresponding binding fragments. (c) Using the data to train ChemPLAN-Net (See Figure 2). (d) FEATURE vectors are extracted from the query protein based on the surface topology of the structure using fpocket and evaluated by ChemPLAN-Net to find potential binding fragments. (e) A list of potential binding fragments is obtained with top predictions ready to be mapped onto query inhibitors.

### Architecture of ChemPLAN-Net

Mapping multiple binding fragments to one query protein environment is a nontrivial computational task, for which there is no general out-of-the-box solution. In order to deal with the multi-label and multi-class problem, we have developed a novel approach using our domain knowledge by creating a well-curated fragment database. Instead of mapping a protein-based input to a fragment-based output, as in traditional end-to-end models, the individual relationship between environments and each fragment in the database is explored through a binding probability descriptor. Figure 2 shows that our model contains two inputs: Firstly, the FEATURE vector is connected to the ResNeXt architecture. Secondly, the Morgan Fin-gerprint of a fragment is concatenated to the output of the ResNeXt and passed through a fully connected layer to a binary classification output, indicating the binding or non-binding label. During the training process, the environment-fragment pairs processed from the PDB files are used with their corresponding binding fragment labels in addition to the artificially generated nonbinding fragments (See Supplementary Note 1). In the testing process, the query environment is run against all fragments in the fragment database, resulting in a list of probabilities that indicate which fragments are most likely to bind.

**FIG. 2.**
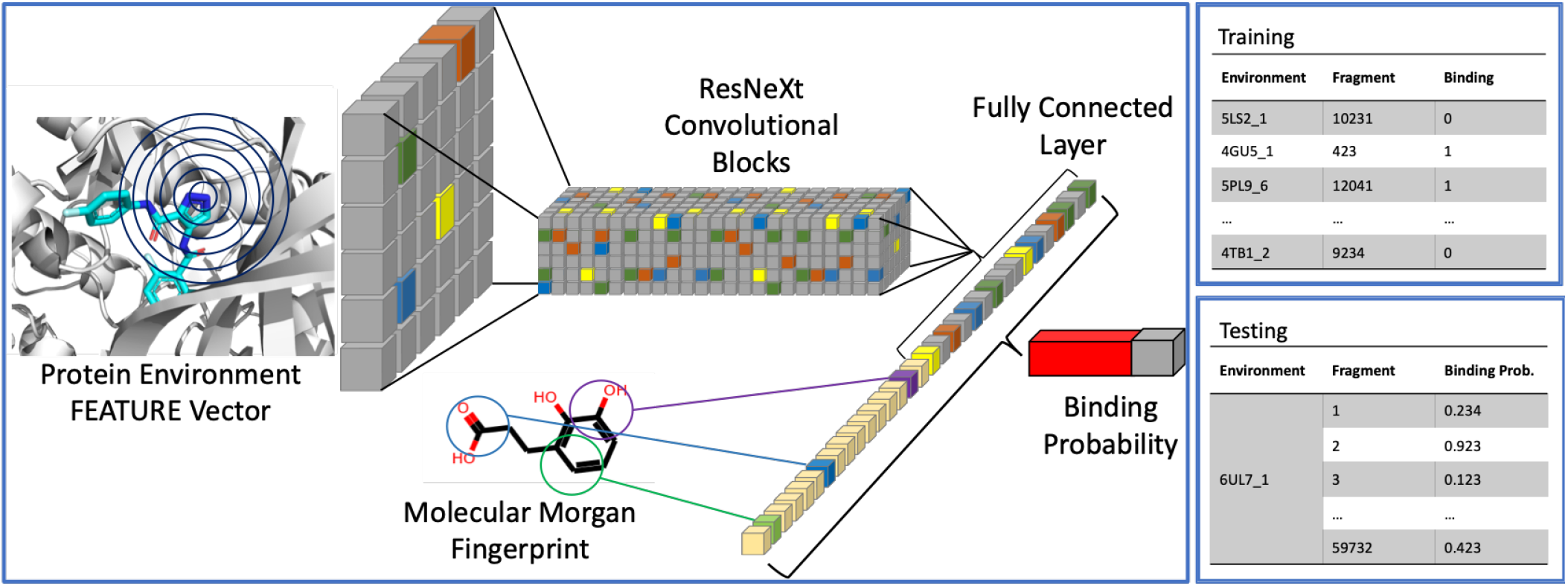
Overview of the ChemPLAN-Net Architecture. The extracted protein environment FEATURE vector is fed into the ResNeXt Convolutional Block, where the higher-order connections are extracted. The fragment is converted into a Morgan Fingerprint and concatenated to the FEATURE vector output. A binding probability is produced for this environment-fragment pair. Training consists of using all the binding and nonbinding environment-fragment pairs extracted from the co-crystal structure and our nonbinding fragment data assumption, respectively. Testing iterates through the 59,732 fragments in the fragment database for every query environment to produce a list of binding probabilities with values closer to one, indicating a higher likelihood of binding.

### Conceptual advancement with respect to the FragFEATURE protocol

We believe that our deep learning method represents a conceptual advancement over the FragFEATURE protocol, because of its phenomenal capability to predict novel ligand fragments that do not exist in the training dataset. It has to be noted that a simple replacement of the nearest neighbor search in the FragFEATURE protocol^11,12^ by a deep learning framework will only result in a method with limited predic-tion power and transferability that is comparable to the old FragFEATURE protocol: i.e., only fragments in the training dataset can be predicted. Furthermore, a direct implementation will impose a major challenge when training the deep learning framework due to the large number of fragments in the training dataset (multi-class problem). The key conceptual advancement of our method is its ability to convert ligand fragments into Morgan Fingerprints and combine them with corresponding FEATURE vectors that describe the chemical environments of binding pockets. This allows our method to implement a powerful deep learning framework that can predict novel ligand fragments that are not present in the training dataset.

To demonstrate the conceptual advancement of ChemPLAN-Net over the FragFEATURE protocol, we designed a control experiment, where we removed 5% of fragment entries from the training dataset (co-crystal structures in the PDB data-bank, see Suppl. Fig. 1A), to assess whether our model could predict the novel fragments that were not present in the training dataset. First, we note that the traditional FragFEATURE protocol is based on a nearest neighbor search that identifies potential binding fragments based on their relevance and frequency in the knowledge base. Therefore, any fragment predictions from the FragFEATURE protocol must originate from this set of fragments. Indeed, the FragFEATURE protocol fails to recover any of the novel fragments, as shown in Suppl. Fig. 1B. However, distinctive from the FragFEATURE protocol, our deep learning method, through the additional use of molecular descriptors, can remarkably predict a significant number of novel fragments that have not been previously featured in the training dataset. We show that our deep learning method was able to correctly identify the majority of the removed fragments ( 70% of novel fragments have been successfully recovered, see Suppl. Fig. 1C). We have also repeated this control experiment with different sets of fragments to exemplify the consistency of our model (Suppl. Fig. 1), and to demonstrate that the remaining fragments contain sufficient chemical information to make accurate predictions for previously unseen fragments across the entire chemical space. These findings highlight the superior learning ability of our deep learning method, which is significantly more transferable and applicable in a new research setting where the use of novel inhibitor fragments may be desired.

### Validation of ChemPLAN-Net by HIV-1 protease

A reoccurring issue with deep learning models is the lack of understanding of the context of the results and their relative significance. To this end, we demonstrate the strength of our model through a careful step-by-step validation to determine its effectiveness, statistical metrics, and potential drawbacks. We approach this by controlling as many variables as possible during training and/or testing and scrutinizing the consequential outcomes of the model. All of the following validation studies are performed on a protease trained model with all entries of HIV-1 protease and sequence homologs removed from the training set. The HIV-1 protease entries are used as the validation set. Controlled validation will be conducted on the over 100 PDB structures of HIV-1 protease, and their over 100 native ligands are considered as the ground truth. Firstly, we will demonstrate the advantage of our deep learning architecture over the random forest (RF) implementation. Secondly, we benchmark ChemPLAN-Net against molecular docking and study the strengths and weaknesses of the prediction algorithm using specific examples. Thirdly, we will show that ChemPLAN-Net has a comparable performance when using ab-initio methods to find binding pocket environments and study the quality of the predictions by quantifying their precision rate. Lastly, we will use experimental in-vitro assay and HTS data to highlight the power of our algorithm beyond structural validation.

### Analysing the fragment prediction performance given known binding environments as demonstrated on HIV-1 protease

Realistic research-setting-like predictions beyond simple binary classification require the selection of binding fragments with the highest probability score. We analyze the predicted fragments and develop a metric on how to quantify the prediction accuracy when compared to ground truth ligands. To this end, the FEATURE vectors of the co-crystal structures of the HIV-1 protease validation dataset are used to predict the binding fragments from the fragment database, which allows us to assume that the binding locations on the protein are known. The protease trained ChemPLAN-Net model (excluding HIV-1 protease) with the relaxed separation parameter at < 25% similarity (See Methods) is used for the analysis. Across the 100 HIV-1 protease PDB structures, over 30 different residues, resulting in 1571 distinct known binding environments, are involved in the binding of the various native ligands. However, the conformation of the protein is very similar amongst the different PDBs, causing the resulting local FEATURE vectors to be numerically very similar. Consequently, the query for this section is only conducted on a subset of (228 out of 1571) known binding environments of HIV-1 protease after removing FEATURE vectors with significant numerical overlap.

The 228 environments are tested against all 59,732 fragments in the fragment database, resulting in a 228 × 59, 732 probability matrix. The fragments with a confidence probability of 0.97 or higher are selected as predicted fragments. This lead to an average of 100 predictions per environment, with 143 unique predictions across all 228 environments (Figure 3d). This indicates that, due to the similarity of the FEATURE vectors in the binding site, the predictions are quite consistent throughout. Furthermore, local deviations most likely originate from the different physicochemical properties of the residues, but considering that all FEATURE vectors are centred on heteroatoms of residues, there is an intrinsic bias against predicting non-polar residues.

**FIG. 3.**
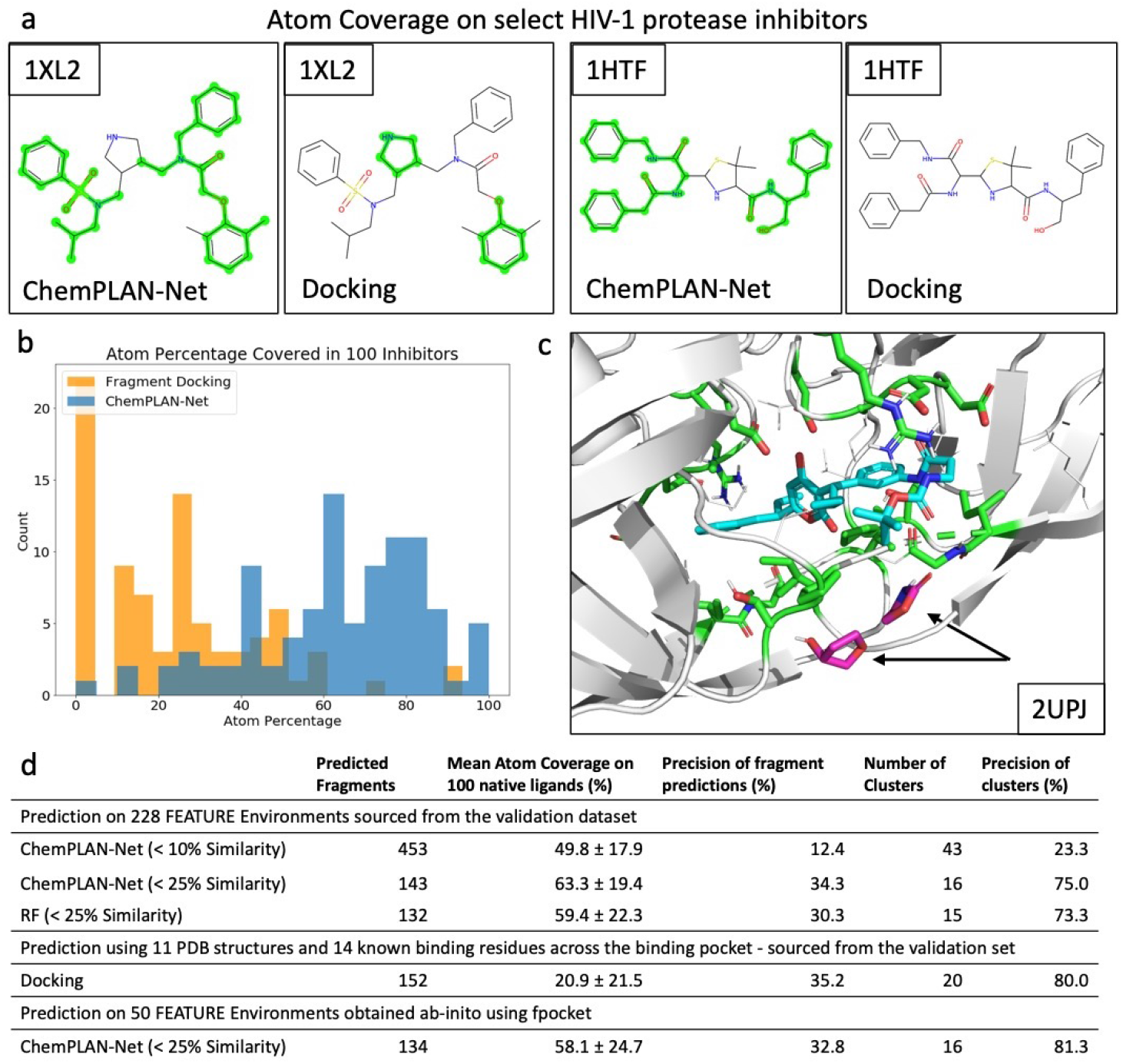
Atom coverage on select HIV-1 protease inhibitors. (a) Side-by-side comparison of atom coverage (green highlight) on two inhibitors with their PDB ID. ChemPLAN-Net can effectively recover non-polar fragments, fragments containing peptide bonds, and is more sensitive to variations containing a benzene substituent, whereas docking predictions are limited to strongly polar small fragments with less regard to their contribution within a larger ligand. (b) Atom coverage across the 100 inhibitors using the top predicted fragments by ChemPLAN-Net and docking (AutoDock Vina). Results show a significantly better coverage for ChemPLAN-Net compared to docking when utilizing a smaller but more diverse fragment set, indicating a higher accuracy with comparable precision. (c) Visualized fragment docking prediction (pink, black arrows) to the residue environments (green) on 2UPJ within an 8 Å grid box. The native ligand is shown for reference (turquoise). All predictions with a negative binding affinity are outside the binding pocket but interact with the protein environments. Docking to binding environments within the pocket leads to highly positive binding affinities, indicating a little binding interaction. (d) Tabular summary of results.

Using ChemPLAN-Net to make targeted predictions poses the challenge of having a very high number of predictions per given query environment. This is an expected result that is already reflected in the training data, where one chemical environment usually binds to multiple fragments that are structurally very similar (differing by one or two atoms). Consequently, instead of attempting to recombine the fragments into a larger molecule, which poses multiple challenges on its own, externally proposed drug molecules are screened against the predicted fragments to find the atom coverage (overlap) in the query drug.

Interestingly, the unique 143 predicted fragments reveal several groups that share a similar structural backbone and chemical features. We evaluate their quality by mapping them back onto the co-crystallized native ligands, as follows: given a native ligand, is there one or more fragments among the predicted fragments (> 0.97 probability) that are a complete substructure of the ligand? If so, what is the percentage of total number of atoms covered on the native ligand by these fragments? (See Methodology for details). The results of this study are displayed in Figure 3d, which shows that among the 100 native ligands, an average of 63% of atoms were covered with a standard deviation of 19%. In practice, this outperformed the RF implementation of the algorithm which has a mean of 59% and standard deviation of 22% using a comparable number of predicted fragments. More significantly, the histogram shows that there were fewer native ligands with a 0% coverage rate, indicating that ChemPLAN-Net was able to reveal at least one crucial fragment that was not predicted by the RF model. Additionally, ChemPLAN-Net predicted slightly more diverse fragments, as shown by the 16 clusters (compared to RF’s 15 clusters) when grouping predicted fragments together based on their similarity. This additional prediction diversity explains the 20 more native ligands at 80+% atom coverage. A detailed discussion of prediction performance in terms of the underlying chemistry can be found in Supplementary Note 2.

### Benchmarking ChemPLAN-Net against molecular docking

We choose molecular docking to perform this benchmark test because both molecular docking and ChemPLAN-Net are structural-based inhibitor prediction methods that primarily use the same input sources, namely structural data. We conduct a thorough analysis using AutoDock Vina^19^ to highlight the advantage that our method has in terms of accuracy and resolution, while also having a faster calculation time. To this end, we select 11 representative HIV-1 main protease structures and perform docking around 14 specified binding environments across the binding pocket using a subset of the 1500 fragments from the fragment database due to the much slower query time. To accelerate the calculations and facilitate direct comparison with ChemPLAN-Net, we ensured that the subset contained 500 known-binding fragments, 500 top predicted fragments by ChemPLAN-Net, and 500 random fragments. We compute the binding affinity of all binding environments against all fragments and limit the search grid box around the specified environment to 8 Å. For every environment, we select the top 30 fragments with the lowest negative binding affinity, resulting in an overall prediction of 152 fragments across the whole binding pocket. Most protein environments inside the pocket are not able to recover any negative binding affinities. This results in an atom coverage ratio of only 21% ± 22% of atoms on the over 100 native ligands from the previous section; however, ChemPLAN-Net successfully achieved 63% ± 19% atom coverage on the native ligands (Fig. 3b). Fig 3a shows that ChemPLAN-Net can effectively recover non-polar fragments, fragments containing peptide bonds, and is more sensitive to variations containing a benzene substituent, whereas docking predictions are limited to strongly polar small fragments with less regard to their contribution within a larger ligand. We attribute this profound advantage to our method’s ability to transfer the binding information gathered from proteins and fragments based on their configurations in the training data. In contrast, docking treats the query fragments as stand-alone molecular entities without considering the context of a potential superstructure ligand. Consequently, docking predictions are located much further from the intended binding sites, which leads to an overall weaker prediction quality. In fact, as shown in Fig. 3c, all predicted fragments with very negative binding affinities are located outside the binding pocket despite being given the ability to bind within the pocket.

### Precision of the model without given binding environments demonstrated on HIV-1 protease

Evaluating the statistical accuracy of the model with respect to the predictions made (i.e., prediction precision) is important to evaluate their significance. The open-ended nature of the problem has to be noted, i.e., the binding fragments that are considered the ground truth are theoretically not an exhaustive list, as they only capture fragments originating from PDB files. In the previous section, the fragment predictions with respect to the native ligands and their atom coverage were quantified. In this section, the predicted fragment space will be explored and a precision metric is defined based on the chemical and structural similarity of the predictions. Furthermore, the methodology used for extracting FEATURE vectors from new query protein crystal structures using the protein’s surface topology has to be validated using a research-like setting.

In order to simulate the research scenario of a less studied novel protein target, only 11 out of the 100 PDB HIV-1 protease structures were used. While the ligands of those selected structures vary in size, the absolute difference is very small with an RMSD between individual structures of less than 0.5 Å. The ligands of those PDB files were removed, and the fpocket software^20^ was run to identify potential binding residue heteroatoms centres. Fpocket uses the surface topology of the protein and a geometry-based Voronoi-tessellation method for the partitioning and identification of cavity-forming residues on both buried and solvent-exposed protein surfaces. The FEATURE vectors are constructed on those 130 identified partially overlapping potential binding environments, of which around 50 are in the known-binding pocket. The other 80 residues are distributed seemingly randomly across the protein. If the binding site is not known, computing the average of the residue predictions across the multiple PDBs will usually show an overlap or have a higher density around the binding pocket.

Each of the 50 environments is tested using ChemPLAN-Net against the approximately 59,732 fragments in the fragment database, and their corresponding probabilities are collected. Using a cutoff probability of > 0.97, an average of 114 predicted fragments are obtained per environment, with 134 unique predicted fragments across the 50 environments in the binding pocket. The strong overlap of predicted fragments between environments is expected as the FEATURE vector of the environments in the binding pocket are in close proximity and are thus very similar. For control, these predictions have been compared to the FEATURE vector of ten of the discarded outlier environments distributed in various locations outside the binding pocket, and a minimal overlap (10% on average) of the predicted fragments was found. The number of predicted fragments, as well as the identified environments, strongly correlate with the predictions made in the previous section when using the known FEATURE vectors obtained from the holo (ligand-bound) structure instead.

The 134 unique predicted fragments can be clustered into 16 chemically and structurally significant clusters based on their Tanimoto Similarity using Agglomerative Clustering,^21^ an unsupervised learning clustering algorithm that is particularly useful for dense fully connected data sets such as a similarity matrix. The number of clusters is chosen by inspection using chemical intuition to separate the fragments accurately with the aim of having about 7-10 fragments per cluster. Representative predicted fragments of six clusters are shown in Supplementary Figure 5. Results show that the fragments are reasonably split as Tanimoto similarity primarily takes the local heteroatom environment types into account before the number of atoms. The individual fragments of the cluster predictions share many similarities within each cluster, such as specific ring backbones, substituent types, chain lengths, and orientation. Further, hierarchical dependencies between clusters can be observed, with some clusters being more similar than others.

Overall, the statistical precision, i.e., the true positive predictions over the total positive predictions is evaluated by computing whether the predicted fragments in the clusters have a substructure in the 100 native ligands of the HIV-1 protease. Results show that 44 fragments out of the 134 predictions can be mapped onto the native ligands, leading to a precision of 32.8%. However, when considering the chemical and structural properties of the predicted fragments, the 44 predicted fragments are shown to belong to 13 out of the 16 predicted clusters. Taking this into account, we can consider an effective precision of 81.3%. For comparison, there are 59,732 fragments in the fragment database with the 100 native ligands mapping to 1600 unique fragments. At best, a random prediction of 130 fragments would achieve a 2.7% precision if individual fragments are considered, and around 8.4% if the fragment space is clustered into chemically meaningful clusters of 50 fragments each.

### Testing ChemPLAN-Net’s inhibitor predictions using in-vitro data without available co-crystal structures of HIV-1 protease

We have demonstrated the ability of ChemPLAN-Net to recover ligand fragments using structural data as validation. To further validate its effectiveness and highlight its usefulness for research-like applications, we introduce a further metric for analysis and compare the predicted fragments against two experimental studies; in-vitro cell assays and high-throughput screening. To match the research-like setting well, we use the 134 fragment predictions made in the previous section from the 50 environments in the binding pocket identified by the ab-initio fpocket method.

We choose seven in-vitro cell studies in both MT-4 and MT-2 cells that measure the inhibition of selected inhibitors on wild-type HIV-1 protease.^22–28^ The studies all report inhibition of various inhibitors in the order of 4-9 pEC50 on wild-type HIV-1 protease, and use a clinically approved HIV-1 protease inhibitor as a baseline for reporting the inhibition constants. Most of these studies are primarily focused on organic synthesis, such that the proposed inhibitors are mutations from the clinically approved inhibitors, with side-chains altered to various degrees. We select the top 5-7 inhibitors from each study that show the highest overall inhibition and structural diversity and end up with 33 total inhibitors. Mapping our predicted fragments for HIV-1 protease shows a large coverage rate for all 33 inhibitors (over 50% coverage), with the best ten inhibitors reporting over 80% coverage (Fig. 4a). We report both the atom coverage, as well as the normalized atom coverage in brackets (when different) that takes into account linear atom structures not present in the fragment database. As in the previous section, we see similar patterns, with some less frequent heteroatoms not being recovered particularly well. For example, one predicted cluster contains the tetrahydrofuran fragment appended to a peptide-like side chain, slightly different than the proposed inhibitors, and therefore does not map back. Generally, no direct correlation between atom coverage and pEC50 values are observed, as many of the chemical and structural properties relevant for protein binding might not be equally relevant for cell penetration or diffusion to the target.

**FIG. 4.**
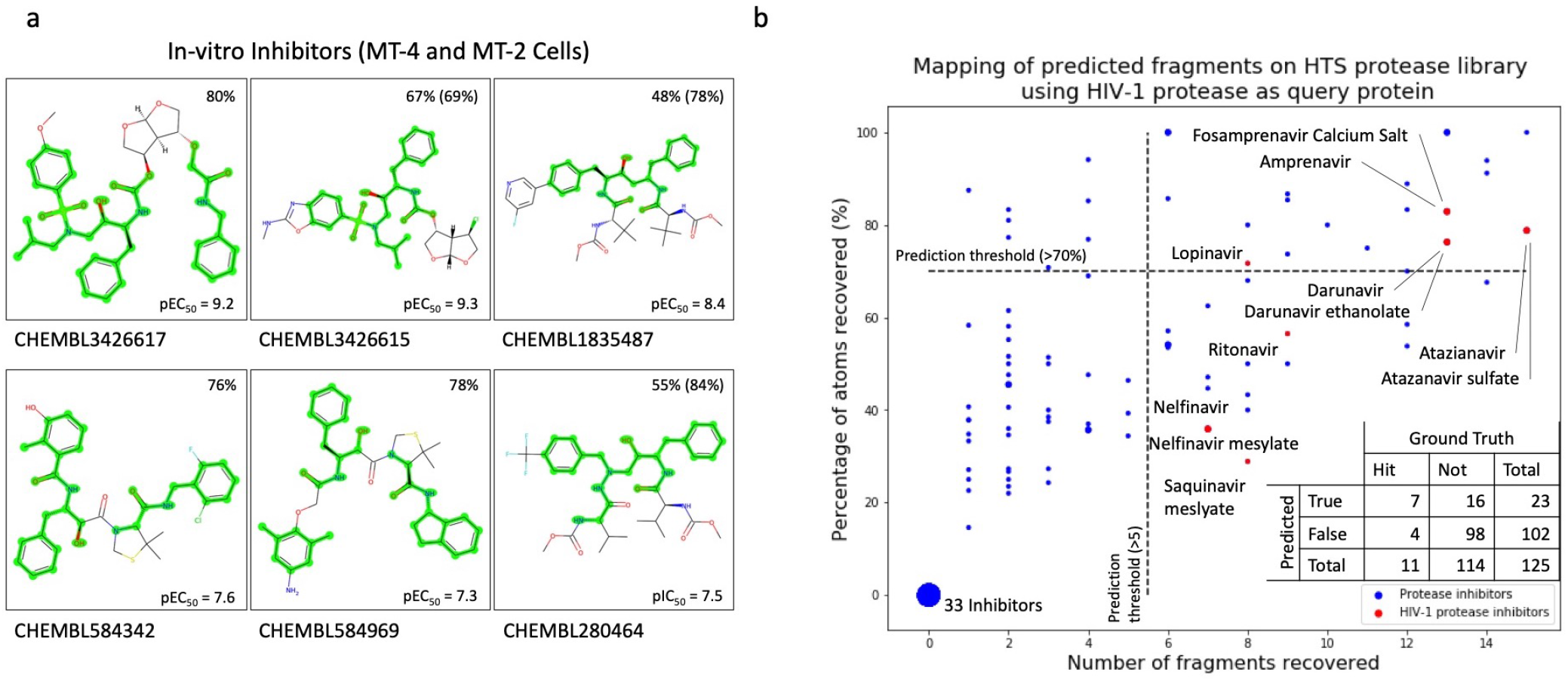
Experimental in-vitro validation. (a) The predicted fragments for HIV-1 protease are mapped back onto in-vitro inhibitors of select assays and atom coverage is reported together with their pEC50 values. Atom coverage in brackets takes into account atoms that are not covered by the fragment database. (b) Predicted fragments for HIV-1 protease are mapped onto 125 protease inhibitors used in a high-throughput screening study, with inhibitors in red being declared as experimental hits. Prediction thresholds for the theoretical predictions are at >70% atom coverage and >5 recovered fragments to compensate for very small compounds or redundant fragment predictions. The table shows the confusion matrix of the predictions.

We select one high-throughput screening study^29^ which measured the inhibition of 130 protease inhibitors on HIV-1 protease, allowing us to benchmark our method against experimentally validated positive and negative results. Figure 4b shows our 134 predicted fragments mapped onto all 125 molecular protease inhibitors and plots the atom coverage percentage of the predicted fragments against the number of fragments recovered per compound. We add the latter condition as some protease inhibitors show very repetitive local structures so that a high atom coverage can be achieved by only recovering one or two very common fragments (i.e., toluene). We define cutoffs for both metrics to define our prediction compounds for inhibition as indicated by the black dotted lines at > 70% atom coverage and >5 number of fragments based on the observations of the previous studies. Results are shown in the top right corner, with 7 out of our 23 predictions showing experimental inhibition, giving a statistical precision of 30.4%. Most other positively predicted inhibitors have a similar backbone and chemical features to the experimental hit compounds. There is a particular high specificity (ratio of true negative over all negatives) value of 86.0%, which indicates the effectiveness of ChemPLAN-Net to predict a small number of prominently featured fragments, rather than a large number of diverse fragments. This is further shown in the bottom left corner, where 33 of the protease inhibitors do not recover any fragments, and consequently show no atom coverage. Other experimentally confirmed inhibitors also score relatively high in the two metrics despite not being featured in the top predictions, indicating that this prediction is better than random. The importance of the second new metric is highlighted, as there is a clear division along the x-axis originating from one or two highly frequent predicted fragments being mapped back onto multiple sites on the inhibitors. In those cases, higher atom coverage did not correspond to a better prediction.

### Further Discussion

The current implementation of ChemPLAN-Net outputs fragment prediction clusters that can be mapped back onto query inhibitors, and their atom coverage is used as an estimate on the usefulness of these inhibitors. It is an advancement of our previous work, NucleicNet,^30^ where we developed a deep learning 7-class classification method to predict the binding preference of RNA constituents on protein surfaces. Rather than solely describing the protein site, like in NucleicNet, we incorporate molecular descriptors of the binding ligand fragments in the form of Morgan Fingerprints for enhanced performance in ChemPLAN-Net. In this study, we focus on targeted drug candidates currently used in ongoing trials where experimental literature is available for validation. However, due to easy software implementations of large databases, screening against already approved drugs in the Drugbank^31^ or potential molecule candidates in the ChEMBL database^32^ is straightforward and allows the prediction of novel inhibitor compounds. Proposed metrics for screening are: the number of predicted fragments covered in those ligands, a diversity score that measures how many different predicted clusters those fragments originate from, and a total number of predicted fragments. This takes advantage of using unseen fragment predictions when evaluating compounds by setting appropriate thresholds for those metrics. In this study, we make little distinction between binding environments within one binding pocket due to their proximity and the limited resolution of the FEATURE framework. This is reflected by the similar fragment predictions made by ChemPLAN-Net for spatially neighbouring FEATURE environments. However, for proteins with larger binding pockets, clear distinctions are observed for different residues. This allows for higher specificity when mapping back to larger compounds, i.e., predictions from different sides of the binding pockets must be mapped back to opposite sides of the ligand. A different approach includes using the predicted fragments from a medicinal chemistry drug-design perspective when optimising lead compound scaffold backbones. For instance, using an existing drug while mutating the side-chains or adding functional groups with the objective to match the predicted fragments onto the structure where appropriate to optimize lead compounds. Naturally, this requires the input and expertise of synthetic chemists to maximise the potential of those predictions. A more challenging approach is the development of an ab-initio algorithm or a generative model that combines predicted fragments to produce chemically and synthetically relevant compounds given a number of predicted fragment inputs. The difficulty is that joining two fragments often requires the overlap of carbon atoms or bonds. Therefore, it is more challenging to implement than simply linking two radical fragments together, as it requires the modification of the input fragments and has to take into account specific desired chemical properties, such as drug likeliness, synthesizability, and solubility.

## CONCLUSION

Fragment-based drug discovery is a versatile field with many approaches over the last decade. Here, we demonstrated the effectiveness of ChemPLAN-Net - an end-to-end pipeline developed to train only on the three-dimensional structures of the PDB protein co-crystal structures - through thorough validation using the well-studied HIV-1 protease. By considering the association of features from binding pocket environments and inhibitor fragments, ChemPLAN-Net supersedes the FragFEATURE protocol in its ability to predict novel fragments that are not found in the training data. This significantly enhances the prediction power in a real-world application as it allows the exploration of novel fragment compounds beyond currently available co-crystal structure data. Learning the associations of the pairwise physicochemical features of proteins and ligands has allowed our method to significantly increase prediction ability at better resolution and accuracy when compared to our molecular docking benchmark. This paper focuses on proteases as the protein family under the assumption that the local binding sites are similar enough to capture relationships between the input space. This framework holds great promise to be trained on other protein family data given this condition. New challenges have to be considered and addressed, such as distinguishing multiple binding pockets, conformation plasticity, and the influence of other chemical species relevant to the inhibition.

## METHODS

### Architecture Overview

The deep learning backbone of ChemPLAN-Net’s residual block architecture was motivated by the successful results of our previous work with NucleicNet.^30^ Here, we use ResNeXt^33^ - a very similar framework with comparable performance, but with the additional benefit of using group convolutions to reduce the number of overall training parameters and thus reduce training complexity. We use a convolutional neural network over a simple feed-forward hidden artificial neural network (ANN) due to the complexity of the input space of 6 × 80 features and to avoid the common pitfalls that come with optimizing ANNs such as computational space-time complexity and the vanishing gradient problem.^34^ Another strength of this implementation is its use of the radial structure of the FEATURE vector, where the six rows of 80 features correlate with the relative distance of those properties to the central atom. Similar to CNN’s use in image classification and its ability to correlate neighbouring pixels and extract underlying meaning in a hierarchical manner, ^35^ the higher-order spatial relationships of properties along the distance to the central atom are explored. The kernel matrix of the ResNeXt blocks operate locally on the input features, allowing it to capture the sequential feature transition across the neighbouring radial layers on the protein binding site in the early layers, and later captures the overarching features in later layers. ChemPLAN-Net uses seven residual blocks with a total of 65 layers, cardinality (group convolution parameter) of eight, and a kernel filter size of 2× 80, 2× 20, and 3× 3 for the first four, subsequent two, and last layer, respectively. The channel size increases sequentially from 1 to 1024 before the resulting high dimensional tensor is reduced using a 3D pooling operation and concatenated to the molecular 1024 bit vector of the Morgan Fingerprint. The final sigmoid layer uses binary-cross-entropy loss to map the output on a probability scale from 0 (non-binding) to 1 (binding). In total, 65 layers are compiled. Hyper-parameter optimisation is conducted, and the model is trained to convergence within 5 epochs at a duration of 16 h on a cluster of 8 Nvidia GeForce GTX 1080 Ti GPUs.

### Sequence Homology and Environment Redundancy

Accurate validation of deep learning models requires careful splitting of training and testing data so that entries of the same protein do not feature in both sets. To this end, protein sequence similarity is taken into account as even proteins differing by a few residues (e.g., through mutations) featuring in both data sets would bias the validation process sufficiently. The fundamental idea of the validation process is to simulate the model’s ability to predict new fragments in a real-world research application, i.e., when a novel query protein has a few similarities to the proteins in the training data. To this end, the BLAST+ tool^36^ is used in the PDB Database to identify all sequence homologs of HIV-1 protease of > 40% and remove their entries from the training data. When curating the training data environment, entries of multiple instances of the same protein structures were often very similar, despite being bound to slightly different ligands. Consequently, to remove redundancy, those environments were merged by concatenating their fragment labels, and duplicate entries were discarded.

### Capturing Physicochemical Environments with FEATURE

Describing the binding environment on the target-protein for machine learning tasks has been essential for successful modelling using deep learning. ^37^ One crucial property of the description vectors is rotational invariance, i.e., the descriptor of the local environments should be insensitive to measurement orientation. In other words, in 3D space, a descriptor that captures physicochemical properties radially is needed, and would ideally be count-based. Fortunately, the FEATURE vector framework has been developed, which operates as follows: Firstly, the space around a functional centre is divided into six concentric spherical shells, each 1.25 Å thick, leading to a total captured space with a radius of 7.5 Å (c.f. benzene diameter is 2.5 Å). Secondly, these shells are each enumerated based on 80 physicochemical properties, including atom and residue type, secondary structure, charge type, etc. (See Supplementary Table 1). Definitions for the functional centres and the properties under consideration are discussed in Ref.. ^12^ By counting the number of these 80 properties over the six concentric spherical shells, each binding environment is described by a 6×80 vector that can be used as an input for ChemPLAN-Net.

### Defining the Fragment Library

The native ligands in the PDB database span over a large chemical space, so that a library of chemical stable fragments were selected as representatives. Chemical intuition was used to choose fragments from the ZINC15 database^38^ based on the following properties: Firstly, they are chemically stable species, thus removing radicals created by “cutting” bonds in half when fragmentising, allowing them to be used for other computational methods (like docking) or purchased for other experiments. Secondly, they should contain aromatic or heterocyclic ring structures (usually with side chains) as they are prominent in many larger drug molecules. Further, using too small fragments like < 5 heavy atom fragments such as carboxylic acids or alcohol groups would create too much noise when fragmentising as they are present in most ligands. Thirdly, limiting the fragments to ≤ 13 heavy atoms in order to maintain relevance as larger fragments tend to appear much less frequently. Some exceptions to those rules are made when chemical relevance has been given precedence for less frequent substituents, such as in the case of nitrile, fluroform, and sulphate groups. A visualisation of the fragment library with a few examples can be found in Supplementary Figure 4. In total, 59,732 purchasable fragments can be derived from ligands deposited in the PDB, which cover 92.6% of atoms on average per ligand. Fragmentising a native ligand for the preparation of training data is done via a superstructure search of all fragments in the fragment database with respect to the native ligand. Consequently, on average, a native ligand ends up with around 20-40 partially overlapping fragments depending on its size and structure.

### a. Atom Coverage

We evaluate the quality of fragment predictions by mapping them back onto the cocrystallized native ligands, as follows: given a native ligand, is there one or more fragments among the predicted fragments that are a complete substructure of the ligand? If so, what is the percentage of total number of atoms covered on the native ligand by these fragments? All fragments contain ring structures and are chemically stable compounds. Therefore, mapping multiple fragments with perfect substructure matches onto the native ligand will lead to atom overlap. I.e. using benzoic acid (C6H5CO2H) as native ligand and toluene (C_6_H_5_CH_3_), benzene (C_6_H_6_) and aniline (C_6_H_5_NH_2_) as predicted fragments would lead to an atom coverage of 7/9 = 77.7%, 6/9 = 66.6% and 0 respectively. The aniline prediction is rejected as it is not a perfect heavy-atom substructure match with benzoic acid. Likewise overall, only the two oxygen atoms in benzoic acid are not featured in the predicted fragments, therefore we assign a atom coverage percentage of 77.7%.

### Morgan Fingerprints and Nonbinding Fragments

Encoding molecules has been an ongoing problem for decades.^39^ Approaches range from simple 3D coordinates of the molecule, over graph-based SMILES^40^ strings, to fingerprint methods that capture and count unique local atom connectivity via a hash table.^41^ While all of them have their merits, fingerprints seem particularly useful in the deep learning framework as their encoded representation has a fixed predetermined length regardless of the size of the molecule, and are rotational invariant as they are constructed on a count-based system. In particular, Morgan Fingerprints are used to convert a 2D chemical structure into a binary 1024 bit-Vector of specified size through a hash table. It has been shown that they retain the chemical features and connectivity of the molecules through the hash table architecture.^41,42^ The metric used to identify the most dissimilar fragments for the nonbinding fragment data generation is Tanimoto similarity - a very popular^43^ metric that is calculated between two binary vectors **a** and **b** of length *k* as follows:

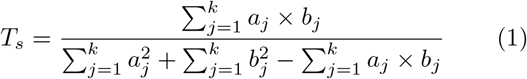

In order to validate this assumption, a separation parameter λ is defined, and three instances of ChemPLAN-Net are trained using the same protein binding data but differing complementary nonbinding fragment data. For the first instance, the nonbinding data is chosen at random amongst the 59,732 different available fragments in the fragment database (random λ). For the second instance, the nonbinding data is selected by implementing a strict high separation λ (< 10% Similarity) where the chosen fragments differ physicochemically the most from their binding fragment counterparts. For the third instance, a relaxed λ (< 25% Similarity) is chosen to obtain nonbinding data that contains fragments both significantly different but also somewhat similar to their binding counterparts.

### Data Description, Training and Testing ChemPLAN-Net

Due to the unconventional architecture of ChemPLAN-Net, the training and testing processes are detailed. The training data set consists of around 3000 protease PDB files with their matching ligands, which have been fragmentised using a superstructure search. The 74,000 binding environments of the proteins are extracted around the native ligand and paired with their binding fragments. On average, one environment binds to around 23 different fragments that often have a backbone overlap. If an environment has more than one binding fragment, the FEATURE vector is duplicated and paired up with the other fragments so that each environment-fragment pair can be treated separately. All 1.7 million binding environment-fragment pairs are assigned a one as a label, indicating that they are binding. After an equal amount of nonbinding fragments are derived for each environment, the nonbinding environment-fragment pairs are assigned a zero as a label, indicating that they are nonbinding. All these environmentfragment-label pairs (minus the HIV-1 protease entries for the validation process) are fed into ChemPLAN-Net for training.

Once the model is trained, two approaches for testing are taken. Firstly, the preliminary validation process (See Validation), where the previously omitted HIV-1 protease environment entries are tested together with their binding fragments, as obtained from their PDB structures. The output is evaluated purely on whether it is correctly categorised into binding and nonbinding. Secondly, for the real prediction, the query PDB files are run through the FragFeature framework to obtain the query FEATURE environment vectors around the binding pockets. These query environments are then sequentially tested against all 59,732 Fragments in the fragment database, and their output is collected as a list of probabilities indicating their binding preferences.

### Comparison Machine Learning Models

Before introducing deep learning, the hypothesis that the combination of FEATURE vectors and Morgan Fingerprints can lead to conclusive results has to be validated. This is achieved twofold; once by using a Support Vector Machine (SVMs) Model^44^ and secondly, a random forest model^45^ on the same training and validation data. SVMs were used for their easy implementation and minimal need for parameter optimization. Linear SVMs are a powerful Machine learning tool but suffer from linearity as well as increasing model size with the number of support vectors increasing drastically with the training data. However, as a first instance, SVMs have a good application as a baseline model. Additionally, a Bagging Classifier^46^ was used as an ensemble approach to enhance the accuracy of the model. It does this by initialising ten instances of the SVMs and aggregating the results. RF Classification models are ensemble learning models comprised of multiple decision trees that are optimised to split the input data space into binary classification labels. In order to accelerate the convergence of the 100 tree initialised model, the maximum depth parameter has been set to 100, reducing the need to purify every single lead in the decision trees, i.e., coarse-graining the input space slightly. RFs have shown to be one of the best performing classification algorithms, often outperforming neural network architectures due to their ability to extrapolate missing data input space and resistance to over-fitting due to their large number of initialised decision trees.^47^

### Data and code availability

The data that support the findings of this study are available from the corresponding author upon reasonable request. The authors declare that all other data supporting the findings of this study are available within the paper and its supplementary information files. The source code for a working version of ChemPLAN-Net is available at https://github.com/masvsuarez/ChemPLAN-Net.

## ACKNOWLEDGEMENTS AND AUTHOR CONTRIBUTIONS

The authors would like to acknowledge the support from the Padma Harilela Endowment Fund to X.H., and the King Abdullah University of Science and Technology (KAUST) Office of Sponsored Research (OSR) under Award No. FCC/1/1976-17, FCC/1/1976-23, FCC/1/1976-26, URF/1/3450-01, URF/1/3412-01, URF/1/4098-01-01, REI/1/0018-01-01, and REI/1/4473-01-01 to X.G.. The authors wish to thank Congmin Yuan, Gu Hanlin and Yu Li for helpful discussions. M.S. designed the framework, implemented the code, and wrote the manuscript. X.G. and X.H. supervised, initiated this study, and wrote the manuscript M.X. and J.H.M.L. extracted FEATURE datasets from PDBs. This research made use of the computing resources of the Supercomputing Laboratory at King Abdullah University of Science. Figure 1 was created by Ivan Gromicho. Scientific Illustrator, Research Communication and Publication Services. Office of the Vice President for Research. King Abdullah University of Science and Technology.

## Notes

### Competing Interest Statement

The authors have declared no competing interest.

