## Supplemental materials for "ChemPLAN-Net: A deep learning framework to find novel inhibitor fragments for proteins"

(Dated: 8 August 2021)

### SUPPLEMENTARY NOTE 1. OPTIMIZATION OF THE NONBINDING DATA PARAMETER ON PREDICTION RESULTS.

Two-class classification networks, such as the binding-nonbinding classifier, are best trained using an equal number of data from both classes. Since verified non-binding data is not available for individual environment-fragment pairs, the nonbinding fragments were selected based on how structurally and physicochemically dissimilar they were from the published binding fragments in the fragment database. That is, fragments that are chemically and structurally very different to a protein environment's set of binding fragments should intuitively not bind. The metric used to identify the most dissimilar fragments is Tanimoto similarity[1] based on the chemical Morgan Fingerprints[2] of the fragments. In order to validate this assumption, a separation parameter $\lambda$ is defined, and three instances of ChemPLAN-Net are trained with different nonbinding fragment data (See Supplementary Fig. 4). For the first instance, the nonbinding data is chosen at random amongst the 59,732 different available fragments of the fragment database (random $\lambda$). For the second instance, the nonbinding data is selected by implementing a strict high separation $\lambda$, where the chosen fragments are the most physicochemically dissimilar from their binding fragment counterparts. For the third instance, a relaxed $\lambda$ is chosen to obtain nonbinding data that contains a mixture of fragments that can be significantly different, structurally, but share some functional similarities to their binding counterparts.

These three trained instances of the ChemPLAN-Net (with different $\lambda$ for the nonbinding data selection) are compared to their equivalent support vector machine (SVM) and RF trained models in their ability to distinguish between binding and nonbinding fragments. This is done by evaluating their statistical performance on this two-class classification task. The expected output of the models given one testing environment-fragment pair is binary - either zero or one, which is achieved in the different frameworks as follows: In ChemPLAN-Net, the final layer converts the probabilities ranging from zero to one into a two-class output using a sigmoid layer at a 0.5 cutoff. SVM constructs and fits support vectors as hyper-planes through the dataset and splits the output into either one of the two classes. Similarly, RF forms multiple sets of decision trees based on the input and classifies the output into one of the two classes.

Results show that increasing the separation parameter increases the prediction accuracy of all three models, as defined by their ability to assign the correct binary label to the HIV-1 protease testing environment-fragment pairs. This shows that our model is able to effectively map an existing higher-order correlation between the molecular and protein input data. Choosing the nonbinding data at random $\lambda$, irrespective of their physicochemical and structural properties, leads to a 50% accuracy in all three models, indicating a random prediction (Supplementary Fig. 4). Using a stricter separation parameter, that only includes nonbinding fragments very dissimilar from the binding fragments ($< 10\%$ similarity), leads to significantly better results with an accuracy of 84.7% and 90.7% for the ChemPLAN-Net and RF model, respectively. At this strict $\lambda$ definition, only very different molecules are assigned to each binding-nonbinding dataset, facilitating the construction of simple decision boundaries across the algorithms. However, the theoretical performance does not reflect equally good results when applied in practice (Fig. 3d) as the top predictions of ChemPLAN-Net are more relevant than those of RF. The use of a more relaxed separation parameter ($< 25\%$ similarity between binding and nonbinding fragments) leads to a lower binary classification accuracy of 80.3% and 82.2% for ChemPLAN-Net and RF, respectively. However, the trained models show a significant improvement over the $< 10\%$ similarity models when recovering the ground truth inhibitors in practice. Therefore, all future validation and testing studies are performed using a $\lambda$ of $< 25\%$ similarity.

The comparable performance of the RF model with respect to ChemPLAN-Net can be explained in terms of the underlying training data distribution of the native ligand fragments. Amongst the protease co-crystal structures, only a third of the 59,732 fragments can be mapped back to native ligands, i.e., are considered binding fragments. Consequently, when selecting nonbinding fragments for

[a)] Electronic mail:
[b)] Electronic mail:

a specific individual binding environment, most choices will originate from the other two-thirds of the fragments that bind to no other environment, given a strict separation parameter. Contrary, for a more relaxed separation parameter, a larger number of nonbinding fragments will include fragments that possibly bind to a different environment, making the training data more complex, and thus more describable by the deep learning architecture.[3] This causes the accuracy performance gap between the RF and ChemPLAN-Net to shrink at $< 25\%$ similarity compared to $\lambda$ at $< 10\%$ similarity.

The real strength of ChemPLAN-Net comes from its ability to output a confidence probability value for the classification of the individual environment-fragment pair when tested. This is explored in the next section when simulating a practical scenario, where every environment is paired with all 59,732 fragments individually and their comparative performance is ranked in a list of probabilities. Using 0.5 probability as the cut off, as the conventional classification algorithm does, is inconvenient as around a third of the fragments in the fragment database would be considered to be binding. Therefore, filtering out the highest-confidence binding fragments (0.97 probability cutoff) is crucial to make a significant prediction to which end the list of ranked probabilities is required.

### SUPPLEMENTARY NOTE 2. DISCUSSION OF THE CHEMPLAN-NET ALGORITHM

By selecting the two best, two worst, and one average performing native ligand, the effectiveness of the ChemPLAN-Net algorithm is discussed. Supplementary Fig. 3a shows the atom coverage in green of the predicted fragments on the individual native ligand, with some predicted fragments overlapping the native ligand. The four-digit alphanumeric symbol corresponds to the PDB ID of the binding protein file of the native ligand.

1HBV has one of the lowest atom coverage as it is composed of many linear fragments that are not included in the fragment database due to redundancy. Most other ligands, containing only short linear side-chains, are captured by the fragment database, as can be seen in the 1HTG or 1NPA example. 1HTG, in particular, shows less atom coverage, as some rare fragments, like the thiazolidine ring in the centre, are less frequent in the training data. The adenosine analogue fragment should have been recovered, but due to the large size of the native ligand, relative contributions of the specific fragment are sometimes not captured. It should be noted that amongst the predicted fragments, there was a structurally very similar adenosine analogue fragment with a slightly different side-chain. 1NPA represents a more typical prediction with most aromatic and heteroaromatic structures being covered as well as their corresponding linkage chains. Independent linkers, such as the alcohol group central to the native ligand, are difficult to recover as they often do not have a corresponding binding environment on the protein side. 2QNQ shows a very good atom coverage partially due to the symmetry of the native ligand, with both the five-ring pyrrolidine and the sulfonic-benzene groups being recovered. Relevant chloride residues in the ortho-position were not recovered; however, other predicted fragments did show negatively charged substituents such as a hydroxy group in the ortho-position. While not fully accurate, the alternative predictions highlight the versatility of ChemPLAN-Net by providing chemically acceptable alternatives in the correct positions. This would allow further studies to investigate possible mutations on specific substituent groups while still maintaining a correct ligand backbone. 1T7K shows good atom coverage on the four residual benzene rings, with the additional central ring being completely covered despite being slightly unorthodox. This native ligand has also been shown in 3D in the co-crystal structure of HIV-1 Protease (PDBID: 1T7K) using the pymol visualisation software[4] with detected ligand-protein interactions highlighted using yellow dash lines (Suppl. Fig. 3b). As can be seen, ChemPLAN-Net does not consider the cyanide group to be a significant contributor to the binding of the ligand as the functional group points outside the binding pocket and is thus not in proximity of a protein residue. Generally, some native ligands will have fragments contributing much less to the binding through weaker electrostatic interactions (such as nonpolar fragments) or will have fragments spatially orientated outside of the binding pocket and are thus significantly underrepresented in the original training data.

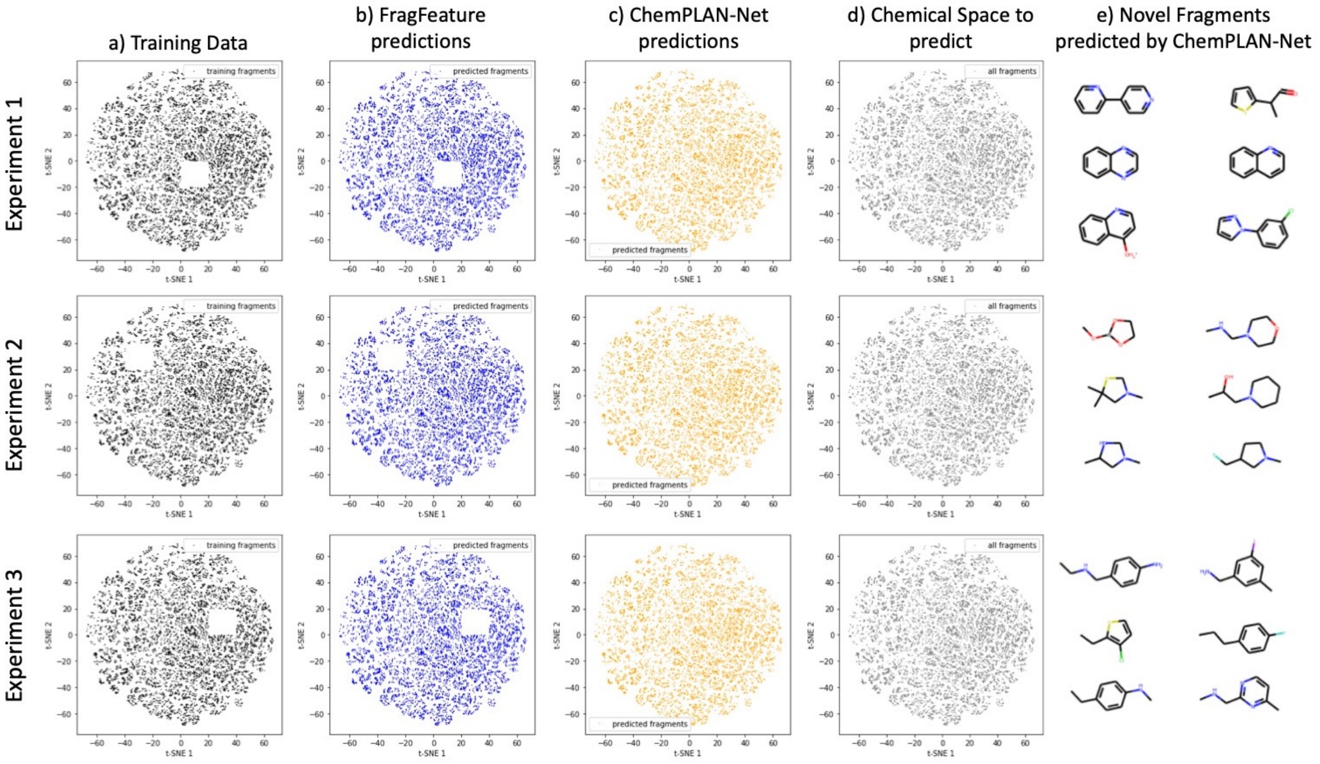


Supplementary Figure 1. 2D projection of the chemical fragment space based on chemical and structural similarity. (a) Select entries (<5% of training data) were removed in each experiment and queries were conducted for environments of various proteins to recover their assigned binding fragments. (b) FragFEATURE predictions are able to recover most environment fragments with the exception of the removed fragments. (c) ChemPLAN-Net is able to recover additional "novel" fragments it has not seen before in the training data due to the implicit learning of molecular fingerprints. (d) We selected a large number of query environments that have binding fragments across the chemical space. (e) Example of novel fragments predicted by ChemPLAN-Net.

### Training Protocol

**Extract Binding Environments and Fragment Labels**
PDB files of the target protein family are obtained from the PDB Database. Binding environment vectors and fragment fingerprints are extracted using the FEATURE package and rdkit respectively. Environment and fragment redundancies are merged and duplicates are removed.

**Creating Nonbinding Fragments**
For each environment the corresponding binding fragments are used to find an equal number of nonbinding fragments that share less than 25% Tanimoto similarity.

**Removing validation protein entries**
All environment/fragment entries of the HIV-1 protease and the 40% homologs are removed.

**Training the model**
The training data is normalised and the environment vectors are duplicated where more than one fragment is binding. A one or zero label for each pair indicates binding or nonbinding respectively. This data is fed into the deep learning framework until model convergence is achieved.

### Validation protocol

**Using removed validation protein entries**
The previously removed environment/fragment entries of the HIV-1 protease and the 40% homologs are used to evaluate the accuracy of the model to classify the pairs as binding or nonbinding using a 0.5 threshold.

### Prediction protocol

**Finding Binding Environments**
Fpocket is used on the query PDB structures to obtain potential binding pockets. FEATURE is applied to vectorise the functional centres to use as query input. Duplicate or almost identical binding environments are removed and binding pockets outside known binding regions are discarded.

**Prediction**
Each environment is iteratively paired up with each fragment of the 59,732 fragment base. This leads to a list of binding probabilities for each fragment with respect to the individual environment.

Supplementary Figure 2. Training, Validation and Prediction protocol of ChemPLAN-Net

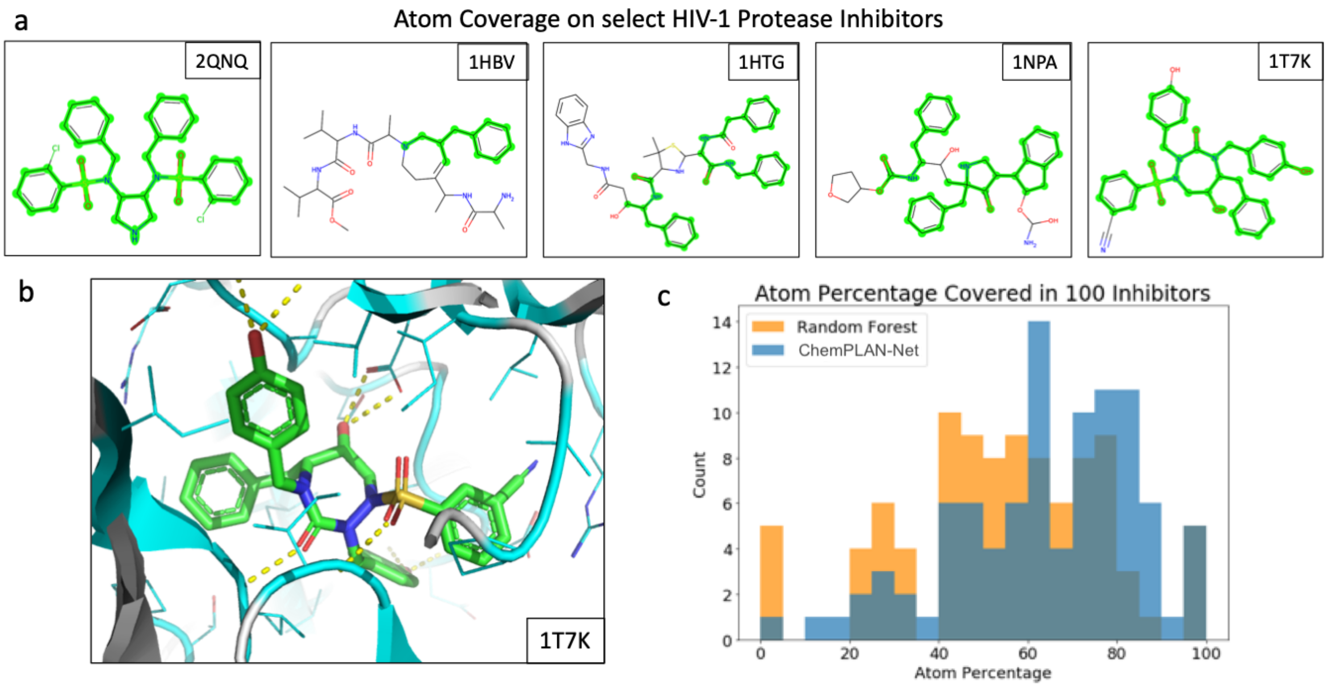


Supplementary Figure 3. Overview of the Validation Results. (a) Atom coverage of predicted fragments (shown in green) highlight the two best, two worst, and one average performing native ligand. (b) The yellow dashed lines show the protein-ligand binding interactions in the 1T7K structure, visualised in pymol. (c) Atom coverage defined by the predicted fragments that cover a substructure of 100 native HIV-1 Protease ligands showing that ChemPLAN-Net (FFN) outperforms random forest when applied in practice. (d) Statistical Results for the validation data predictions using the known binding FEATURE vectors from the validation dataset and the ab-inito FEATURE vectors when sourced through fpocket.

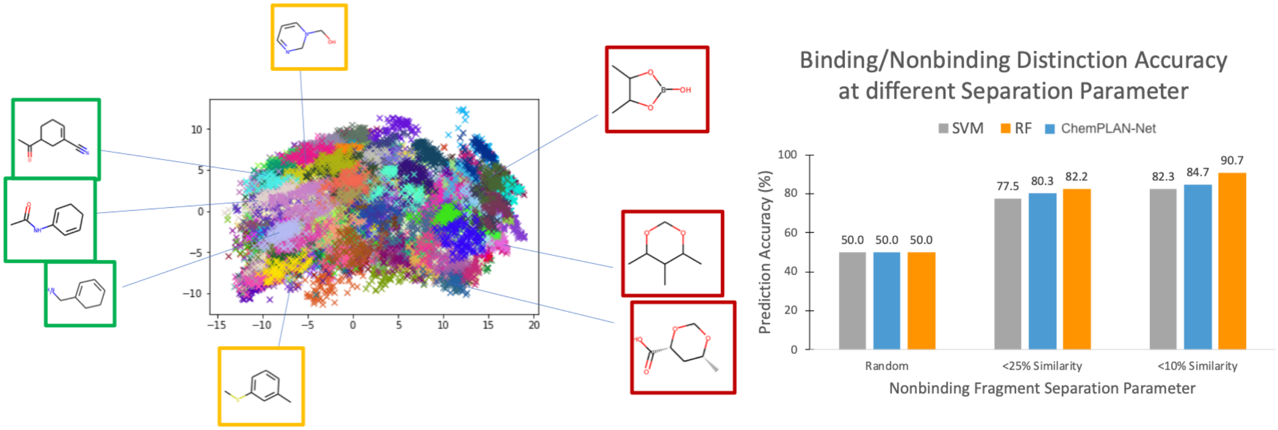


Supplementary Figure 4. (left) Visualisation of Tanimoto similarity in the molecular fragment space. The first two principal components of the Principal Component Analysis of the similarity matrix are plotted against each other. K-means clustering with 500 centres was used to aid visualisation of similar fragments with lower euclidean distance corresponding to a higher similarity. A strict separation parameter ($< 10\%$ Similarity) would split the binding and non binding data into green and red fragments, whereas a relaxed separation parameter ($< 25\%$ Similarity) could include orange fragments in the nonbinding dataset. (right) Accuracy of distinguishing binding vs. nonbinding classification at 0.5 probability threshold for the validation data. Random forest outperforms FNN and SVM using this theoretical metric.

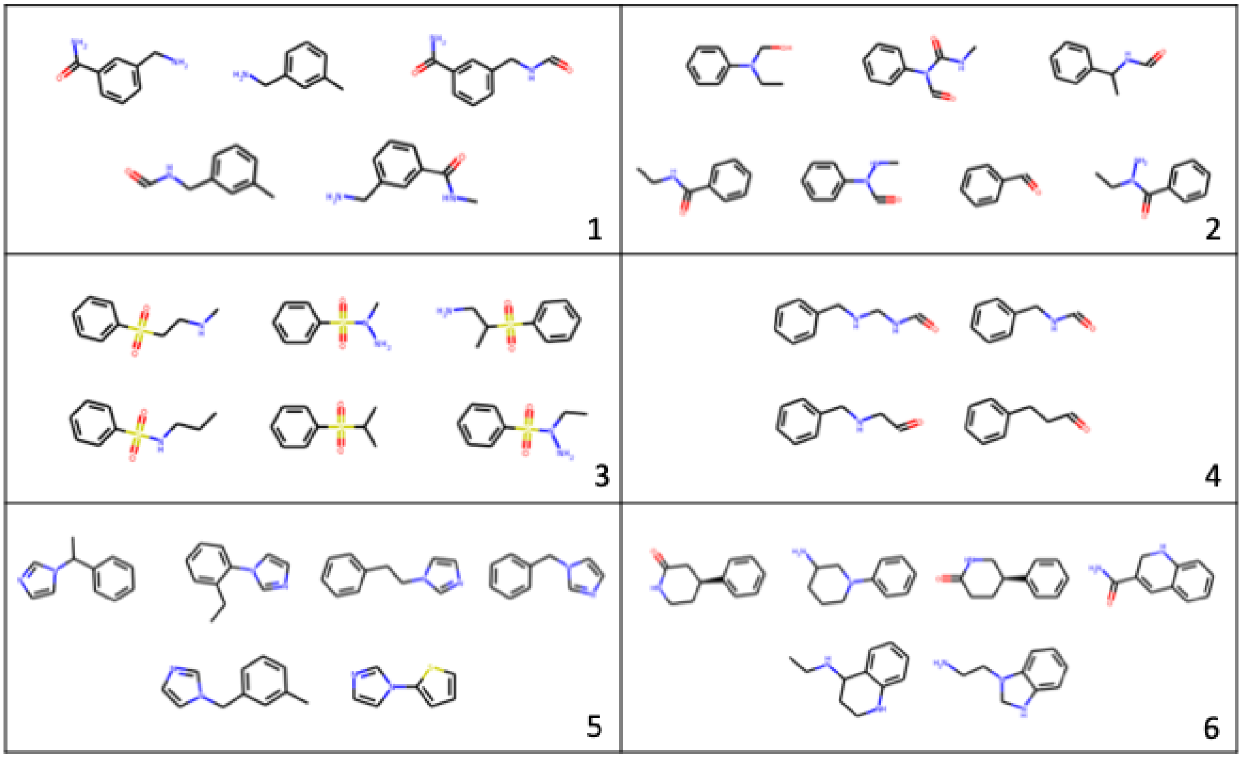


Supplementary Figure 5. 6 representative prediction fragment clusters for HIV-1 Protease inhibitors. ChemPLAN-Net predicts around 134 Fragments for the binding pocket. These predictions are clustered into chemically and structurally meaningful groups reducing the variability of predictions significantly.

| Atom-based | Residue-based | Secondary-structure-based |
|---|---|---|
| ATOM-TYPE-IS-C | RESIDUE_NAME_IS_ALA | SECONDARY_STRUCTURE1_IS_3HELIX |
| ATOM-TYPE-IS-CT | RESIDUE_NAME_IS_ARG | SECONDARY_STRUCTURE1_IS_4HELIX |
| ATOM-TYPE-IS-Ca | RESIDUE_NAME_IS_ASN | SECONDARY_STRUCTURE1_IS_5HELIX |
| ATOM-TYPE-IS-N | RESIDUE_NAME_IS_ASP | SECONDARY_STRUCTURE1_IS_BRIDGE |
| ATOM-TYPE-IS-N2 | RESIDUE_NAME_IS_CYS | SECONDARY_STRUCTURE1_IS_STRAND |
| ATOM-TYPE-IS-N3 | RESIDUE_NAME_IS_GLN | SECONDARY_STRUCTURE1_IS_TURN |
| ATOM-TYPE-IS-Na | RESIDUE_NAME_IS_GLU | SECONDARY_STRUCTURE1_IS_BEND |
| ATOM-TYPE-IS-O | RESIDUE_NAME_IS_GLY | SECONDARY_STRUCTURE1_IS_COIL |
| ATOM-TYPE-IS-O2 | RESIDUE_NAME_IS_HIS | SECONDARY_STRUCTURE1_IS_HET |
| ATOM-TYPE-IS-OH | RESIDUE_NAME_IS_ILE | SECONDARY_STRUCTURE1_IS_UNKNOWN |
| ATOM-TYPE-IS-S | RESIDUE_NAME_IS_LEU | SECONDARY_STRUCTURE2_IS_HELIX |
| ATOM-TYPE-IS-SH | RESIDUE_NAME_IS_LYS | SECONDARY_STRUCTURE2_IS_BETA |
| ATOM-TYPE-IS-OTHER | RESIDUE_NAME_IS_MET | SECONDARY_STRUCTURE2_IS_COIL |
| ATOM-NAME-IS-ANY | RESIDUE_NAME_IS_PHE | SECONDARY_STRUCTURE2_IS_HET |
| ATOM-NAME-IS-C | RESIDUE_NAME_IS_PRO | SECONDARY_STRUCTURE2_IS_UNKNOWN |
| ATOM-NAME-IS-N | RESIDUE_NAME_IS_SER | |
| ATOM-NAME-IS-O | RESIDUE_NAME_IS_THR | |
| ATOM-NAME-IS-S | RESIDUE_NAME_IS_TRP | |
| ATOM-NAME-IS-OTHER | RESIDUE_NAME_IS_TYR | |
| HYDROXYL | RESIDUE_NAME_IS_VAL | |
| AMIDE | RESIDUE_NAME_IS_HOH | |
| AMINE | RESIDUE_NAME_IS_OTHER | |
| CARBONYL | CLASS1_IS_HYDROPHOBIC | |
| RING-SYSTEM | CLASS1_IS_CHARGED | |
| PEPTIDE | CLASS1_IS_POLAR | |
| | CLASS1_IS_UNKNOWN | |
| | CLASS2_IS_NONPOLAR | |
| | CLASS2_IS_POLAR | |
| | CLASS2_IS_BASIC | |
| | CLASS2_IS_ACIDIC | |
| | CLASS2_IS_UNKNOWN | |
| | PARTIAL-CHARGE | |
| | VDW-VOLUME | |
| | CHARGE | |
| | CHARGE-WITH-HIS | |
| | NEG-CHARGE | |
| | POS-CHARGE | |
| | HYDROPHOBICITY | |
| | MOBILITY | |
| | SOLVENT-ACCESSIBILITY | |

Supplementary Table 1. Eighty chemical properties accounted by FEATURE vectors
